# An assessment of the human Sortilin1 protein network, its expression and targetability using small molecules

**DOI:** 10.1101/2023.04.13.536697

**Authors:** Arun HS Kumar

**Affiliations:** Stemcology, School of Veterinary Medicine, University College Dublin, Belfield, Dublin-04, Ireland

**Keywords:** Sortilin1, fibrosis, microcalcification, unstable plaque, lycopene

## Abstract

**Background:** Sortilin1 (SORT1) is a ubiquitously expressed transporter involved in sorting or clearing proteins and is pathologically linked to tissue fibrosis and calcification. Targeting SORT1 may have potential clinical efficacy in controlling or reversing cardiovascular fibrosis and/or calcification. Hence this study assessed the protein-protein network of human SORT1 and its targetability using small molecule nutra/pharmaceuticals.

**Material and methods:** Network proteins of SORT1 in homo sapiens was identified using the String database, and the affinity of the protein-protein interaction of this network was analysed using Chimera software. The tissue specific expression profile of SORT1 was evaluated and assessed for enrichment in different cell types including the immune cells. A library of in-house small molecules and currently used therapeutics for cardiovascular diseases were screened using AutoDock vina to assess targetability of human SORT1. Concentration affinity (CA) ratio of the small molecules was estimated to assess the clinical feasibility of targeting SORT1.

**Results:** IGF2R, NTRK2, GRN and GGA1 were identified as high affinity interaction networks of SORT1. Of these high affinity interactions, IGF2R and GRN can be considered as relevant networks in regulating tissue fibrosis or microcalcification process due to their influence on T-cell activation, inflammation, wound repair, and tissue remodelling process. The tissue cell type enrichment indicated major expression of SORT1 in adipocytes, specialised epithelial cells, monocytes, cardiomyocytes, and thyroid glandular cells. The binding pocket analysis of human SORT1 showed twelve potential drug interaction sites with varying binding score (0.86 to 5.83) and probability of interaction (0.004 to 0.304). Five of the drug interaction sites were observed to be targetable at therapeutically feasible concentration of the small molecules evaluated. Empagliflozin, sitagliptin and lycopene showed superior affinity and CA ratio compared to established inhibitors of SORT1.

**Conclusion:** IGF2R and GRN are relevant networks of SORT1 regulating tissue fibrosis or microcalcification process. SORT1 can be targeted using currently approved small molecule therapeutics (empagliflozin and sitagliptin) or widely used nutraceutical (Lycopene) which should be evaluated in a randomised clinical trial to assess the efficacy to reduce cardiac/vascular microcalcification process.

## Introduction

Sortilin1 (SORT1) is a ubiquitously expressed membrane glycoprotein which functions as sorting receptor in the Golgi apparatus and a clearance receptor on the cells membrane which is vital to functional regulation of cell homeostasis.^[1-4]^ In its role as a sorting receptor, it transports proteins from Golgi apparatus to their intended target site using the endosomes/lysosomes as a cargo, based on a pH based dissociation kinetics.^[3, 4]^ Among the various transporter functions of SORT1, its ability to scavenge extracellular lipoprotein lipase^[5]^ and promote mineralisation of extracellular matrix,^[6]^ contribute to tissue fibrosis, calcification, and foam cell formation.^[7, 8]^ SORT1 seems to favour intracellular accumulation of lipids and triglycerides by suppression and/or degradation of adipogenesis factors and lipid efflux transporters.^[3, 7]^ The expression of SORT1 is highly enriched in adipocytes^[9, 10]^ and smooth muscle cells,^[7, 11]^ which together with its contribution to tissue fibrosis and calcification makes it a vital target in the development of unstable atherosclerotic plaques (UAP) and vascular calcification.

Increased expression of SORT1 is observed in calcified arteries of humans.^[7, 8]^ Consistent with this SORT1 knockout mice show reduced arterial calcification without impacting bone mineral density.^[7, 8]^ This suggest a specific role of SORT1 in pathological microcalcification process without disturbing the physiological bone mineralisation. SORT1 is reported to transport alkaline phosphatase (ALP) to extracellular vesicles (EV)^[12]^ and trigger mineralisation of extracellular matrix (specifically collagen), which leads to functional compromise of otherwise an elastic tissue.^[6, 7]^ The role of SORT1 in microcalcification can collaterally contribute to formation of unstable atherosclerotic plaques (UAP) through the synergistic effects of SORT1 on secreting proinflammatory cytokines (IL6 and IFN gamma) from macrophages.^[13, 14]^ Inflammation and microcalcification are hallmarks of UAP to which SORT1 seems to have a direct contributory role, independent from its effects on lipoprotein metabolism.^[7, 13]^ Interestingly the contribution of SORT1 to UAP is also independent of plasma cholesterol levels.^[7, 13]^ The microcalcification promotion role of SORT1 is not limited to vascular smooth muscles, as recently SORT1 was reported to promote aortic valve fibrosis and calcification following mechanical injury.^[8]^ It appears to me that following tissue injury the transporter function of SORT1 facilitates extracellular matrix deposition (EV dependent and independent mechanisms) which is further reinforced by ALP induced calcification to minimise the impact of the tissue injury. During this calcification process if the endogenous pool of progenitor cells are not efficient to replace the damaged tissue, then the calcified tissue will persist by further mineralisation. There seems to be a transient window during which the activity of SORT1 can be pharmacologically interfered to promote tissue regeneration by either endogenous or exogenous progenitor cells. Hence to address the pharmacological interference of SORT1, this study assessed its protein network, expression profile in human tissues, and its targetability using small molecules.

## Materials and Methods

The network protein analysis of human SORT1 was conducted as reported before using the STRING Database (https://string-db.org), to observe its functional protein-protein interactions.^[15, 16]^ The protein networks were identified and reviewed in the UniProt database and the most resolved protein structure based on the full length of protein and atomic resolution was selected for analysis in this study. The PDB code or AlphaFold structure of the selected protein was used to import the protein structure onto the Chimera software and the number of hydrogen bonds (H-bond) formed between SORT1 and its associated network proteins at 10 Armstrong (10A) distance was evaluated. A heatmap of the number of H-bonds formed between SORT1 and its associated network proteins was generated to identify the high affinity interactions.

The protein expression profile of SORT1 in various human tissues were text mined from the Human Protein Atlas (https://www.proteinatlas.org) database together with the data on enrichment of SORT1 in various tissue cell types. The SORT1 mRNA expression pattern in human immune cells was also assessed from this database. The binding sites of human SORT1 were identified using the PrankWeb: Ligand Binding Site Prediction tool (https://prankweb.cz/). To assess the targetability of SORT1 by small molecules, in this study molecular docking of selected drugs approved for therapeutics of cardiovascular disease were evaluated against AlphaFold structure of human SORT1 using AutoDock vina 1.2.0. Briefly, the AlphaFold structure of SORT1 was optimised for molecular docking in the AutoDock MGL tools. The isomeric SMILES sequence of the selected small molecules were obtained from the PubChem database and were converted into the PDB format using the Chimera software.^[17, 18]^ The PDB structures of small molecules were further processed in the AutoDock MGL tools for molecular docking as reported previously.^[15-19]^ SORT1 AlphaFold structure was also screened against our in-house compound library to identify a high affinity inhibitor. AF38469 and SB203580 which are reported to be orally bioavailable SORT1 inhibitor were used as reference compounds.^[3, 7-9]^ To assess the targetability potential of SORT1 by the small molecules, the concentration affinity ratio (CA ratio) for each of the small molecules was calculated as described before.^[17, 19]^

## Results

The network protein analysis of SORT1 in humans showed high affinity interactions with IGF2R, NTRK2, GRN and GGA1 (Figure 1). IGF2R is a cation-independent mannose-6-phosphate receptor, which facilitates transport of phosphorylated lysosomal enzymes from the Golgi complex/cell surface to lysosomes and also positively regulated T-cell coactivation.^[20, 21]^ NTRK2 is a neurotrophic factor receptor tyrosine kinase which supports development and maturation of nervous system, including establishing nerve-nerve or nerve-glial cell synapses.^[22, 23]^ GRN is a secreted protein which modulates inflammation, wound repair, and tissue remodelling through cytokine like mechanisms influencing fibroblasts and endothelial cells.^[24, 25]^ GGA1 is a binding protein which facilitates a two-way protein sorting and trafficking between the trans-Golgi network (TGN) and endosomes.^[26]^

**Figure 1:**
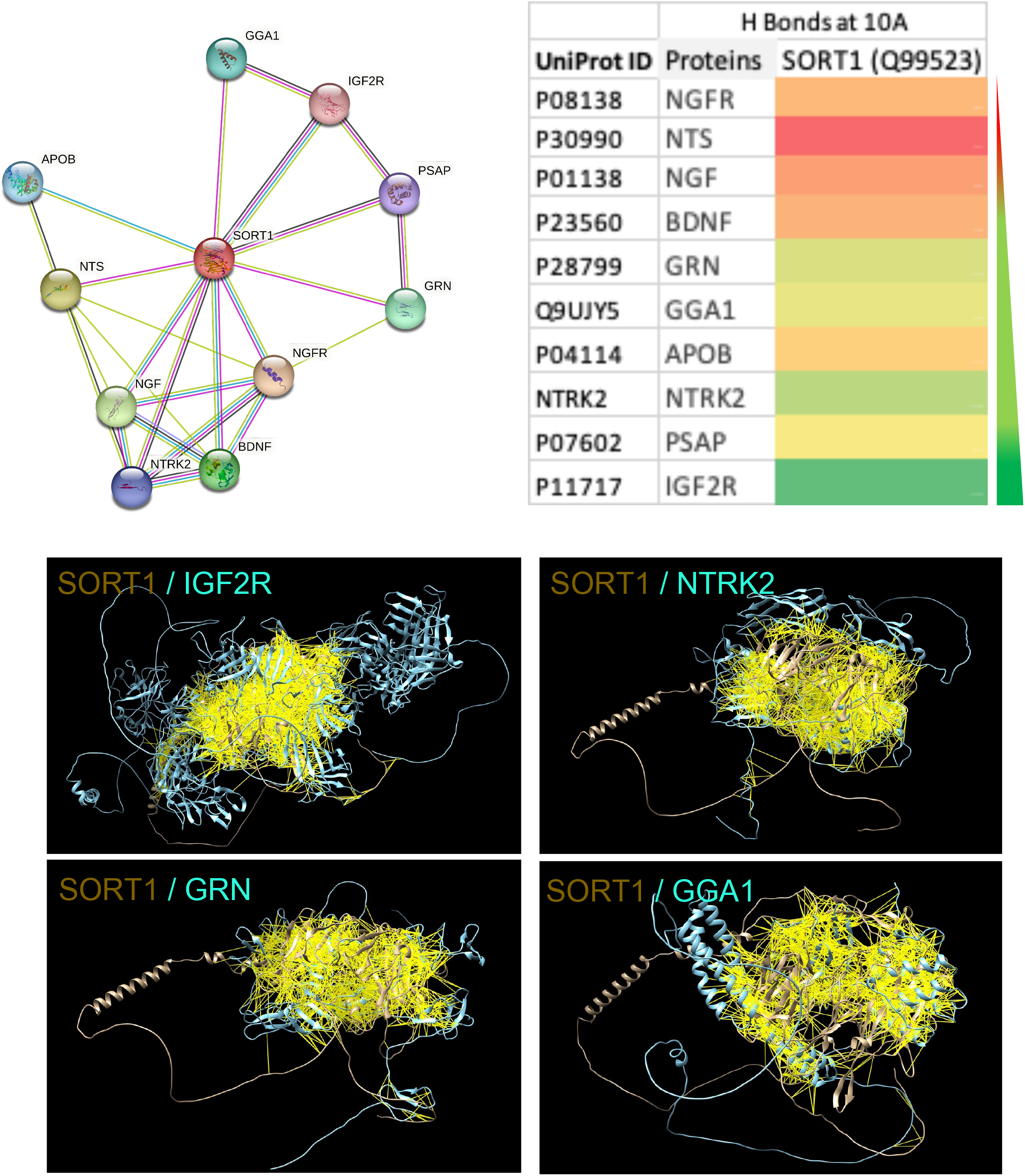
Affinity of human Sortilin1 (SORT1; UniProt ID Q99523) to its network proteins. Network proteins of SORT1 from the string database is shown. The network proteins are represented using their gene code and UniProt ID. Heatmap showing the degree of interaction (number of hydrogen bonds (H Bonds)) between each of the network protein and SORT1 at 10 Armstrong (10A) bond distance. Scale: green to red = high to low. The high affinity network proteins (IGF2R, NTRK2, GRN, and GGA1) in cyan are show interacting (yellow lines represent the H Bonds) with human SORT1 (brown).

The analysis of SORT1 protein expression in human tissues suggested major presence in brain, kidneys, skin, colon, placenta, endocrine glands and testis (epididymis) (figure 2). The tissue cell type enrichment of this data suggested major expression in adipocytes, specialised epithelial cells, cardiomyocytes and thyroid glandular cells (figure 2). The analysis of SORT1 mRNA expression in immune cells indicated significantly higher expression in monocyte subsets (especially nonclassical monocytes) (figure 2).

**Figure 2:**
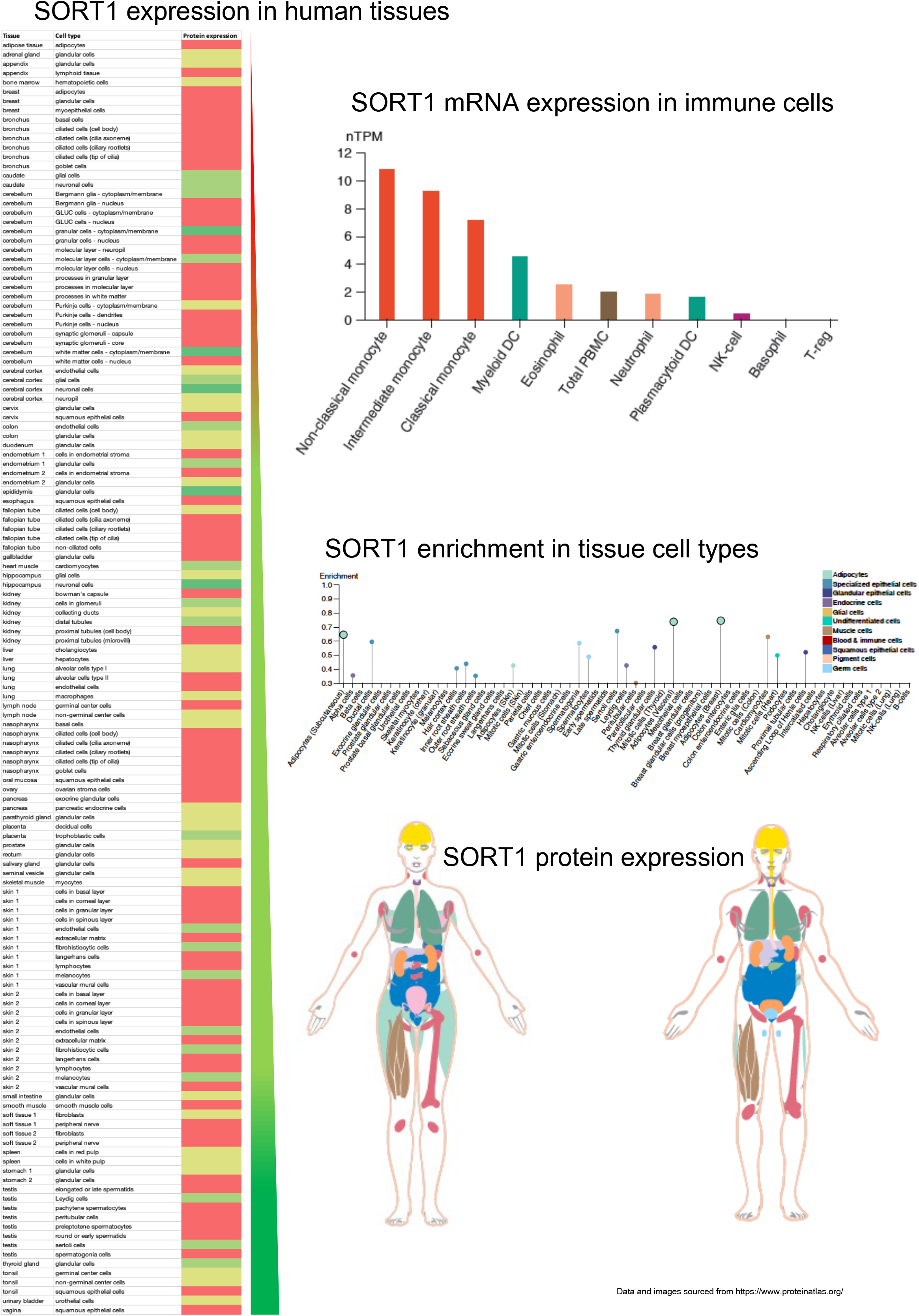
Expression profile of Sortilin1 (SORT1) in human tissues. The quantitative protein expression of SORT1 in various human tissue and cell types is shown as a heat map (Scale: green to red = high to low). The mRNA expression of SORT1 in human immune cells is shown as bar graph (The expression profile is categorised based on the nTPM ranging from 0 to 12 and colour coded based on cell category type). The cell type enrichment (Scale 0 to 1) of SORT1 protein in human tissues is shown as a vertical line graph and colour coded based on cell category type. The gender specific protein expression of SORT1 in various human tissue is shown qualitatively using images.

The binding pocket analysis of human SORT1 showed twelve potential drug interaction sites with varying binding score (0.86 to 5.83) and probability of interaction (0.004 to 0.304) (figure 3). The top binding pocket consisted of 11 amino acids (A432 A631 A634 A635 A636 A637 A667 A669 A671 A672 A718), 48 SAS points and 25 surface atoms with following dimensions (x, y, and z: -29.6892, 3.1138, and -12.6381). The second binding pocket had a score of 4.6 and consisted of 17 amino acids (A341 A377 A386 A387 A388 A435 A674 A675 A676 A677 A678 A679 A680 A681 A691 A697 A698), 66 SAS points and 33 surface atoms with following dimensions (x, y, and z: -31.2511, -5.5778, and 0.9165). The rest of the binding pockets had a score of <3 and probability of interaction of < 0.092 and hence are not described.

**Figure 3:**
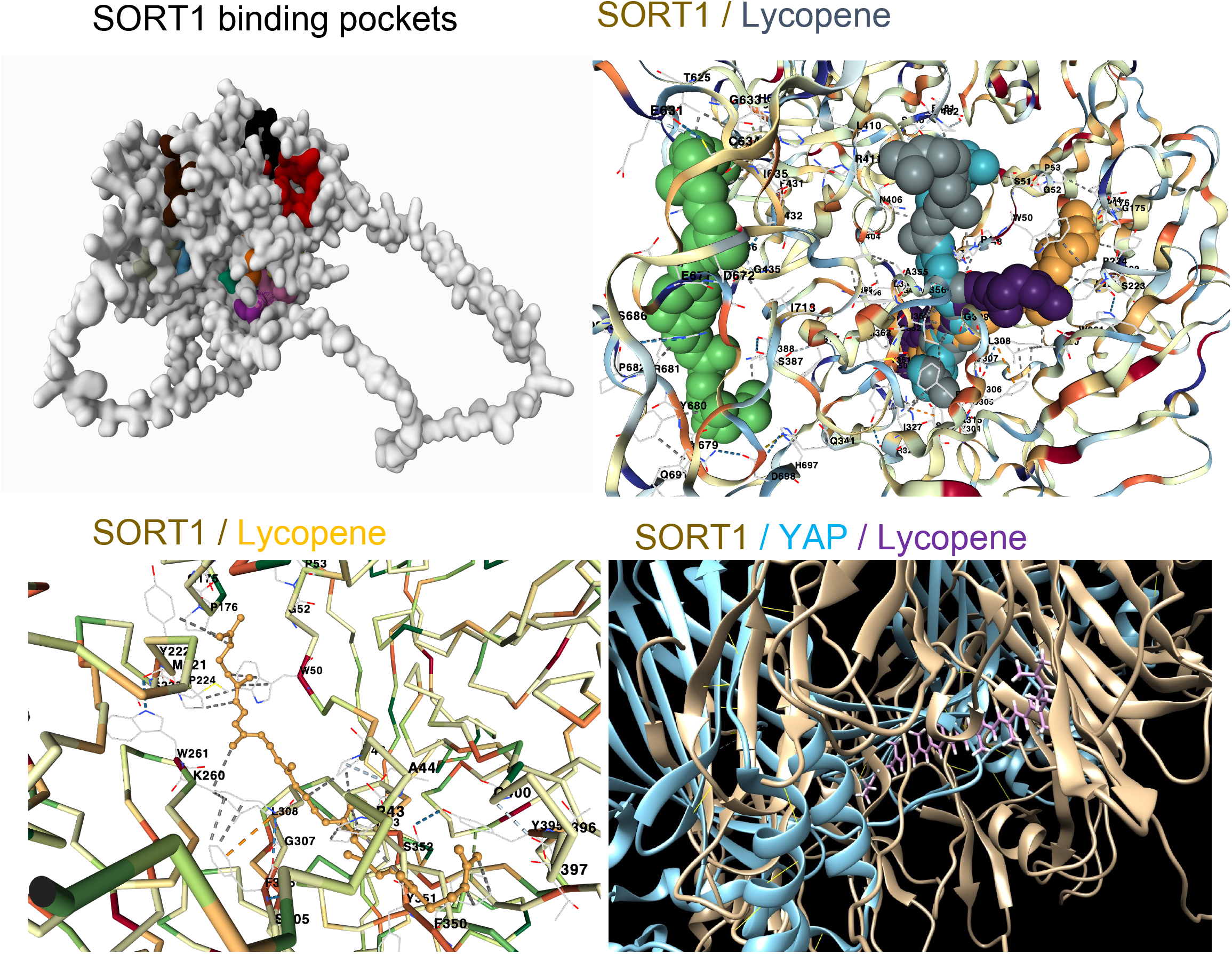
Targetability of human Sortilin1 (SORT1). The twelve binding pockets of SORT1 identified are colour coded and represented in 2D (the relative area of the colour indicates the binding score which ranged from 0.86 to 5.83). The molecular docking of lycopene (ball and stick model) with human SORT1 with its five interaction sites are shown. The bottom left panel shows the highest affinity interaction between lycopene (stick model) with human SORT1. The bottom right panel shows the interaction sites of lycopene (purple stick model) between human SORT1 (brown colour) and yes-associated protein 1 (YAP; cyan colour).

The screening of in-house ligand library against human SORT1 showed high affinity of lycopene against five of the SORT1 binding pockets (Table 1, figure 3). In a recent study SORT1 was reported to contribute to calcification of human aortic valves involving MAPK and YAP pathways which drive trans differentiation of functional cells to myofibroblastic-osteogenic phenotype.^[8]^ Lycopene was observed to bind at several SORT1-YAP interaction sites (figure 3), suggesting its potential to interfere with SORT1-YAP signalling. The potential of various cardiovascular therapeutics to target SORT1 was also analysed (figure 4). Empagliflozin and sitagliptin showed therapeutically feasible affinity and CA ratio against SORT1 similar to that of lycopene (figure 4). Among the small molecules tested bisoprolol had the highest affinity against SORT1, however considering its very low CA ratio, the feasibility to achieve therapeutic safety is unlikely. Two of the orally bioavailable SORT1 inhibitors reported in the literature^[3, 8]^ i.e., SB203580 (3.24 µM) and AF38469 (638.76 µM) showed affinity against SORT1 which was inferior to lycopene (1.23 µM) (figure 4).

**Table 1:** Molecular docking of Lycopene with human SORT1

| Binding Pocket ID | Binding energy (kcal/mol) | Cavity volume (Å<sup>3</sup>) | Centre (x, y, z) | Docking size (x, y, z) | Amino acid sequence |
| --- | --- | --- | --- | --- | --- |
| C2 | -8.2 | 1820 | 5, 7, 3 | 41, 41, 41 | PRO43 ALA44 PRO46 TRP50 GLY52 PRO53 ILE174 GLY175 PRO176 MET221 TYR222 SER223 PRO224 LYS260 TRP261 SER305 PHE306 GLY307 LEU308 PHE350 TYR351 SER352 ILE353 TYR395 THR396 THR397 GLY400 |
| C4 | -8.1 | 666 | 3, 2, -6 |  | PRO46 LEU47 PRO48 ARG49 SER51 TYR304 SER305 PHE306 GLY307 LEU308 GLY309 PHE314 SER316 ARG325 ILE327 PHE350 TYR351 SER352 ILE353 LEU354 ALA355 ALA356 MET363 ASN406 THR408 SER480 GLU481 PRO482 |
| C5 | -8.1 | 445 | -3, -4, 22 |  | PRO43 ALA44 PRO46 LEU47 ARG49 TRP50 PRO224 GLN225 LYS260 TRP261 GLY262 SER263 GLY307 LEU308 GLY309 GLY310 ARG325 GLN347 GLU348 GLN349 PHE350 TYR351 SER352 ILE353 TYR395 THR397 THR398 GLY400 |
| C1 | -7.9 | 3278 | -13, 0, 0 |  | PRO46 LEU47 PRO48 ARG49 TRP50 LYS260 SER305 PHE306 GLY307 LEU308 GLY309 PHE314 ALA315 SER316 ILE327 TYR351 SER352 ILE353 LEU354 ALA355 ALA356 MET363 PHE404 ASN406 THR408 LEU410 ARG411 PRO482 |
| C3 | -7.0 | 1039 | -32, 1, -7 |  | GLN341 PHE377 SER387 LYS388 PHE431 ASP432 GLY435 ARG436 TYR513 HIS623 THR625 GLU631 GLY633 CYS634 ILE635 GLU671 ASP672 TYR679 TYR680 ARG681 PRO682 ASP685 SER686 GLN691 HIS697 ASP698 ILE718 |

**Figure 4:**
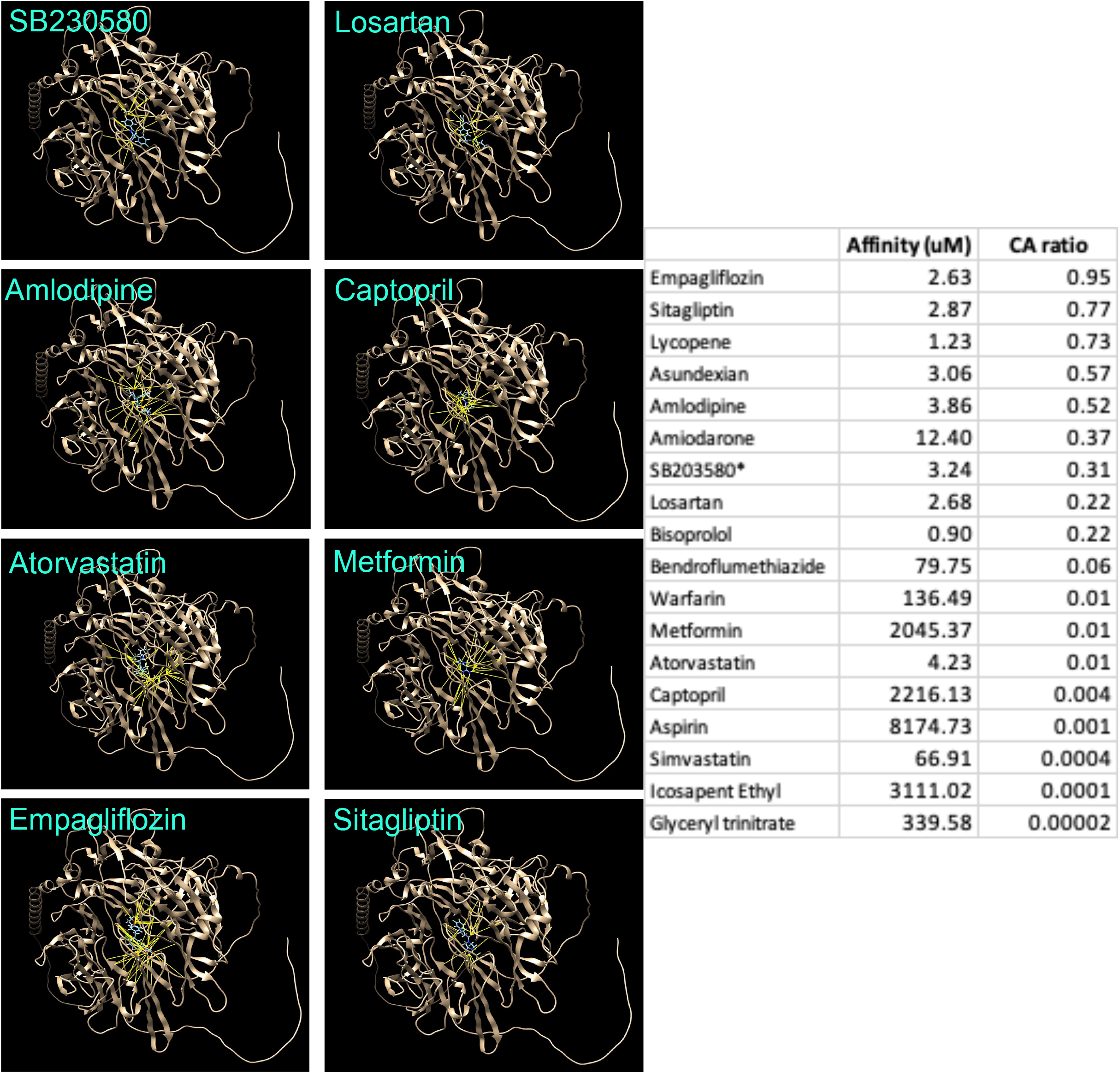
Affinity of small molecules with human Sortilin1 (SORT1). The affinity and concentration-affinity (CA) ratio value of various small molecules against SORT1 is summarised in the table. The images show the degree of interaction (number of hydrogen bonds; yellow lines) and the interaction sites between the selected small molecules and SORT1 (brown colour).

## Discussion

SORT1 is reported to be the driver of microcalcification process associated with cardio-valvular and cardiovascular diseases independent of abnormalities with lipid metabolism and plasma cholesterol levels.^[1, 7, 8, 27]^ Hence this study analysed targetability of SORT1 and has identified new/therapeutic small molecules (Lycopene, empagliflozin and sitagliptin) with affinity to SORT1 at therapeutically feasible range which can be tested for efficacy in a clinical trial.

The major network proteins of SORT1 identified in this study seem to vastly support the transporter function of SORT1. However the SORT1-GRN network is of specific interest within the context of microcalcification process. The GRN dependent modulation of inflammation, wound repair, and tissue remodelling process is synergistic with the SORT1 facilitated extracellular matrix deposition reinforced by ALP induced tissue calcification.^[24, 25, 28]^ A recent study has reported the role of MAPK and YAP pathways in SORT1 induced aortic valve calcification following mechanical injury.^[8]^ While it is possible that biological response to acutely induced mechanical injury may differ from subacute/chronic injury classically seen with cardio-valvular and cardiovascular diseases in humans. The increased expression of SORT1 observed in calcified arteries of humans^[7, 29, 30]^ does suggest that some overlap exists in biological response to acute, subacute and chronic injury to tissues. This concept was validated by a preclinical study simulating acute mechanical tissue injury^[8]^ and pharmacological interventions demonstrating the efficacy of orally bioavailable SORT1 inhibitors in reducing the incidence of microcalcification process.^[7, 8, 31, 32]^ The comparison of the affinity of these SORT1 inhibitors with lycopene, empagliflozin and sitagliptin in this study suggest the potential superior efficacy outcomes from using these small molecules, which will be interesting to be evaluated in randomised clinical trials. The therapeutic use of empagliflozin^[33, 34]^ and sitagliptin^[35, 36]^ in patients who are at higher risk of developing tissue calcification should allow rapid testing of its clinical efficacy in reducing the incidence of calcified aortic valves and/or unstable plaques. Further as lycopene is a generally regarded as safe nutraceutical,^[37, 38]^ a coformulation of lycopene/empagliflozin or lycopene/sitagliptin can also be evaluated. Concerns are also expressed over achieving therapeutic concentration of lycopene in circulation.^[39, 40]^ However the large volume of distribution of lycopene (150 to 1400 litres or 2-19 l/kg) together with its high lipo-solubility suggests that it is extensively distributed into peripheral tissues.^[39, 40]^ Additionally population studies have shown serum lycopene concentrations of >0.7 µM, suggesting SORT1 targetable concentration (CA ratio ∼ = 1) is achievable. In addition to the SORT1 inhibitors other allosteric approachs such as small molecules which can interfere with SORT1-YAP or SORT1-GRN signalling may also be developed as therapeutics for clinical management of calcified valves/arteries.

The expression of SORT1 in cardiomyocytes although is consistent with its pathological role in valvular calcification,^[8, 41]^ its lower expression in vascular smooth muscles contradicts its reported role in arterial calcification. In this context the contributory role of non-classical monocytes (myeloid cells) to formation of viable vascular smooth muscles can be a mechanism by which SORT1 leads to vascular classification.^[42, 43]^ The constant recycling of vascular smooth muscle cells by myeloid cells is an essential process in retaining optimal vascular physiology and the role of SORT1 in this process is unclear. However following mechanical injury to arteries, the myeloid cells promote remodelling of vascular smooth muscles under an inflammatory milieu and paraphs SORT1 plays an active role in this process.^[42, 43]^ The highest expression profile of SORT1 mRNA in the monocyte subpopulation among all immune cells together with its expression in calcified vascular tissue supports its role in pathological vascular remodelling process eventually leading to microcalcification. SORT1 is highly expressed in adipocytes which is consistent with several studies linking the disturbed lipid metabolism with valvular and arterial calcification process.^[3, 44, 45]^ However some preclinical studies have also reported lipid and cholesterol independent mechanisms of SORT1 in inducing tissue calcification.^[41, 46-49]^ The lipid independent role of SORT1 on tissue calcification is of specific interest in the context of UAP, as none of the therapeutics currently used for lipid management are shown to reduce the risk associated with UAP vascular pathology.^[50-52]^ Hence, targeting of SORT1 together with or without lipid management therapeutics can prove to be beneficial in improving vascular physiology by reducing microcalcification process. The potential to target SORT1 using currently approved small molecule therapeutics (empagliflozin and sitagliptin) or widely used nutraceutical (Lycopene) identified in this study is particularly promising and should be evaluated in a randomised clinical trial to assess the efficacy to reduce cardiac/vascular microcalcification process.

## Acknowledgements

Research support from University College Dublin-Seed funding/Output Based Research Support Scheme (R19862, 2019), Royal Society-UK (IES\R2\181067, 2018) and Stemcology (STGY2917, 2022) is acknowledged.

## References

1. Kjolby, M., Nielsen, M. S. and Petersen, C. M. Sortilin, encoded by the cardiovascular risk gene SORT1, and its suggested functions in cardiovascular disease. Current atherosclerosis reports, 2015, 17, 1–9.

2. Ouyang, S., Jia, B., Xie, W., Yang, J. and Lv, Y. Mechanism underlying the regulation of sortilin expression and its trafficking function. Journal of Cellular Physiology, 2020, 235, 8958–71.

3. Su, X. and Peng, D. New insight into sortilin in controlling lipid metabolism and the risk of atherogenesis. Biological Reviews, 2020, 95, 232–43.

4. Mitok, K. A., Keller, M. P. and Attie, A. D. Sorting through the extensive and confusing roles of sortilin in metabolic disease. Journal of Lipid Research, 2022, 63.

5. Nielsen, M. S., Jacobsen, C., Olivecrona, G., Gliemann, J. and Petersen, C. M. Sortilin/neurotensin receptor-3 binds and mediates degradation of lipoprotein lipase. Journal of Biological Chemistry, 1999, 274, 8832–36.

6. Maeda, S., et al. Sortilin is upregulated during osteoblastic differentiation of mesenchymal stem cells and promotes extracellular matrix mineralization. Journal of cellular physiology, 2002, 193, 73–79.

7. Goettsch, C., et al. Sortilin mediates vascular calcification via its recruitment into extracellular vesicles. The Journal of clinical investigation, 2016, 126, 1323–36.

8. Iqbal, F., et al. Sortilin enhances fibrosis and calcification in aortic valve disease by inducing interstitial cell heterogeneity. European Heart Journal, 2023.

9. Morris, N. J., et al. Sortilin is the major 110-kDa protein in GLUT4 vesicles from adipocytes. Journal of Biological Chemistry, 1998, 273, 3582–87.

10. Kaddai, V., et al. Involvement of TNF-α in abnormal adipocyte and muscle sortilin expression in obese mice and humans. Diabetologia, 2009, 52, 932–40.

11. Kapustin, A. and Shanahan, C. Emerging roles for vascular smooth muscle cell exosomes in calcification and coagulation. The Journal of physiology, 2016, 594, 2905–14.

12. Itoh, S., Mizuno, K., Aikawa, M. and Aikawa, E. Dimerization of sortilin regulates its trafficking to extracellular vesicles. Journal of Biological Chemistry, 2018, 293, 4532–44.

13. Mortensen, M. B., et al. Targeting sortilin in immune cells reduces proinflammatory cytokines and atherosclerosis. The Journal of clinical investigation, 2014, 124, 5317–22.

14. Talbot, H., et al. Regulatory roles of sortilin and SorLA in immune-related processes. Frontiers in Pharmacology, 2019, 9, 1507.

15. Manchukonda, B. and Kumar, A. H. Network profiling of hepatocellular carcinoma targets for evidence based pharmacological approach to improve clinical efficacy. Biology, Engineering, Medicine and Science Reports, 2022, 8(1), 11–15.

16. Khosravi, Z., Kaliaperumal, C. and Kumar, A. Analysing the role of SERPINE1 network in the pathogenesis of human glioblastoma. bioRxiv, 2022.

17. Kumar, A. H. Pharmacological targets of Asundexian relevant to its therapeutic efficacy in treating cardiovascular diseases. Biology, Engineering, Medicine and Science Reports, 2022, 8, 24–27.

18. Kumar, A. PTPRC, KDM5C, GABBR1 and HDAC1 are the major targets of valproic acid in regulation of its anticonvulsant pharmacological effects. Biology, Engineering, Medicine and Science Reports, 2022, 8(2), 28–32..

19. Kumar, A. H. Comparative pharmacology of direct oral anticoagulants and vitamin K antagonist. Biology, Engineering, Medicine and Science Reports, 2022, 8(2), 16–23.

20. Yang, G., et al. Insulin-like growth factor 2 enhances regulatory T-cell functions and suppresses food allergy in an experimental model. Journal of allergy and clinical immunology, 2014, 133, 1702-08. e5.

21. Braulke, T. Type-2 IGF receptor: a multi-ligand binding protein. Hormone and metabolic research, 1999, 31, 242–46.

22. Juhasz, G., et al. The CREB1-BDNF-NTRK2 pathway in depression: multiple gene-cognition-environment interactions. Biological psychiatry, 2011, 69, 762–71.

23. Gray, J., et al. Functional characterization of human NTRK2 mutations identified in patients with severe early-onset obesity. International journal of obesity, 2007, 31, 359–64.

24. Jian, J., Konopka, J. and Liu, C. Insights into the role of progranulin in immunity, infection, and inflammation. Journal of leukocyte biology, 2013, 93, 199–208.

25. He, Z., Ong, C. H., Halper, J. and Bateman, A. Progranulin is a mediator of the wound response. Nature medicine, 2003, 9, 225–29.

26. Wahle, T., et al. GGA proteins regulate retrograde transport of BACE1 from endosomes to the trans-Golgi network. Molecular and Cellular Neuroscience, 2005, 29, 453–61.

27. Arvind, P., Nair, J., Jambunathan, S., Kakkar, V. V. and Shanker, J. CELSR2– PSRC1–SORT1 gene expression and association with coronary artery disease and plasma lipid levels in an Asian Indian cohort. Journal of cardiology, 2014, 64, 339–46.

28. Gass, J., et al. Progranulin regulates neuronal outgrowth independent of sortilin. Molecular neurodegeneration, 2012, 7, 1–13.

29. Ryu, J., Ahn, Y., Kook, H. and Kim, Y.-K. The roles of non-coding RNAs in vascular calcification and opportunities as therapeutic targets. Pharmacology & Therapeutics, 2021, 218, 107675.

30. O’Donnell, C. J., et al. Genome-wide association study for coronary artery calcification with follow-up in myocardial infarction. Circulation, 2011, 124, 2855–64.

31. Rogers, M. A. and Aikawa, E. Cardiovascular calcification: artificial intelligence and big data accelerate mechanistic discovery. Nature Reviews Cardiology, 2019, 16, 261–74.

32. Proudfoot, D., et al. Apoptosis regulates human vascular calcification in vitro: evidence for initiation of vascular calcification by apoptotic bodies. Circulation research, 2000, 87, 1055–62.

33. Wanner, C., et al. Empagliflozin and clinical outcomes in patients with type 2 diabetes mellitus, established cardiovascular disease, and chronic kidney disease. Circulation, 2018, 137, 119–29.

34. Kohler, S., Zeller, C., Iliev, H. and Kaspers, S. Safety and tolerability of empagliflozin in patients with type 2 diabetes: pooled analysis of phase I–III clinical trials. Advances in therapy, 2017, 34, 1707–26.

35. Scott, L. J. Sitagliptin: a review in type 2 diabetes. Drugs, 2017, 77, 209–24.

36. Engel, S. S., Round, E., Golm, G. T., Kaufman, K. D. and Goldstein, B. J. Safety and tolerability of sitagliptin in type 2 diabetes: pooled analysis of 25 clinical studies. Diabetes Therapy, 2013, 4, 119–45.

37. Joshi, B., et al. Therapeutic and medicinal uses of lycopene: A systematic review. 2020.

38. Holzapfel, N. P., et al. The potential role of lycopene for the prevention and therapy of prostate cancer: from molecular mechanisms to clinical evidence. International journal of molecular sciences, 2013, 14, 14620–46.

39. Hsieh, M.-J., et al. Cardiovascular disease and possible ways in which lycopene acts as an efficient cardio-protectant against different cardiovascular risk factors. Molecules, 2022, 27, 3235.

40. Moia, V. M., et al. Lycopene used as anti-inflammatory nanodrug for the treatment of rheumathoid arthritis: animal assay, pharmacokinetics, ABC transporter and tissue deposition. Colloids and Surfaces B: Biointerfaces, 2020, 188, 110814.

41. Goettsch, C., Kjolby, M. and Aikawa, E. Sortilin and its multiple roles in cardiovascular and metabolic diseases. Arteriosclerosis, thrombosis, and vascular biology, 2018, 38, 19–25.

42. Kumar, A. H., et al. Role of CX3CR1 receptor in monocyte/macrophage driven neovascularization. PLoS One, 2013, 8, e57230.

43. Kumar, A. H., et al. Bone marrow-derived CX3CR1 progenitors contribute to neointimal smooth muscle cells via fractalkine CX3CR1 interaction. FASEB J, 2010, 24, 81–92.

44. Conlon, D. M. Role of sortilin in lipid metabolism. Current opinion in lipidology, 2019, 30, 198–204.

45. Gao, A., et al. Implications of sortilin in lipid metabolism and lipid disorder diseases. DNA and Cell Biology, 2017, 36, 1050–61.

46. Hu, D., Yang, Y. and Peng, D.-q. Increased sortilin and its independent effect on circulating proprotein convertase subtilisin/kexin type 9 (PCSK9) in statin-naive patients with coronary artery disease. International journal of cardiology, 2017, 227, 61–65.

47. Zhong, L.-y., et al. Sortilin: a novel regulator in lipid metabolism and atherogenesis. Clinica Chimica Acta, 2016, 460, 11–17.

48. Lu, H., Sheppard, M. and Daugherty, A. Calcification in atherosclerotic lesions. Current opinion in lipidology, 2016, 27, 543.

49. Jones, G. T., et al. A sequence variant associated with sortilin-1 (SORT1) on 1p13. 3 is independently associated with abdominal aortic aneurysm. Human molecular genetics, 2013, 22, 2941–47.

50. Migdalski, A. and Jawien, A. New insight into biology, molecular diagnostics and treatment options of unstable carotid atherosclerotic plaque: a narrative review. Annals of Translational Medicine, 2021, 9.

51. Skagen, K., Skjelland, M., Zamani, M. and Russell, D. Unstable carotid artery plaque: new insights and controversies in diagnostics and treatment. Croatian Medical Journal, 2016, 57, 311–20.

52. Makris, G. C., Lavida, A., Nicolaides, A. N. and Geroulakos, G. The effect of statins on carotid plaque morphology: a LDL-associated action or one more pleiotropic effect of statins? Atherosclerosis, 2010, 213, 8–20.

